# Interlinked Bi-stable Switches Govern the Cell Fate Commitment of Embryonic Stem Cells

**DOI:** 10.1101/2023.08.30.555552

**Authors:** Amitava Giri, Sandip Kar

## Abstract

Cell fate decision-making events of embryonic stem (ES) cells are often governed by bi-stable switch-like steady state dynamics of two key transcription factors Oct4 and Nanog due to mutual positive feedback regulation. Intriguingly, for the differentiation of ES cells to either extraembryonic trophectoderm (TE) and primitive endoderm (PrE) lineages, Oct4 maintains either low (TE) or high PrE) expression levels relative to a moderate level in ES cells, while Nanog exhibits high expression in ES state and low expression in both the differentiated states. The dynamical origin for such kind of disparate steady-state level maintenance of Oct4 and Nanog remains elusive. Herein, we demonstrate that such complex steady-state dynamics can be hypothesized and explained by two different bistable switches interconnected either in a stepwise (Oct4) or in a mushroom-like (Nanog) manner. Our hypothesis qualitatively reconciles various experimental observations and elucidates how different feedback and feed-forward motifs orchestrate the extraembryonic development and stemness maintenance of ES cells. Importantly, the model predicts strategies to optimize the dynamics of self-renewal and differentiation of embryonic stem cells that may have therapeutic relevance in the future.

## Introduction

The embryonic development in the mammalian system happens by coordinating precise decision-making at each stage of embryo development^1,2^. Thus, it offers great insights into how different feedback and feed-forward motifs govern these decision-making processes at the single-cell level during early embryonic development^3,4^. Mouse embryonic stem (ES) cells are mainly derived from the Inner cell mass (ICM) of the mouse blastocyst^5–9^. ES cells are pluripotent in nature. They can self-renew themselves, and pluripotency of ES cells is maintained during self-renewal by preventing differentiation and promoting proliferation^10,11^. However, in the presence of appropriate internal and external cues, ES cells can differentiate into the two extraembryonic either trophectoderm (TE) or Primitive endoderm PrE lineages^12–16^. Experimental studies have shown that key intracellular transcription factors, such as Nanog, Oct4, and Sox2 intricately control the cellular fate of ES at the molecular level^17–20^. However, these transcription factors intricately maintain the expression levels of each other by involving in a set of feedback and feed-forward network motifs to orchestrate the cell fate decision-making of ES cells^21,22^. The question remains, how the dynamical events related to these cell-fate decision-making events are governed by these three transcription factors? Importantly, can we alter these cell fate decision-making events to our advantage by perturbing the transcriptional network regulating the differentiation dynamics of ES cells?

Intriguingly, the expression levels of these key transcription factors dictate the fate commitment choice of ES cells^18,21–24^. By engineering ES cell lines (ZHBTc4 and ZHBTc6) where the Oct4 expression level can be varied externally^25^, Niwa et al. have shown that the precise level of Oct4 governs the three distinct cell fates of ES cells. They have observed that a critical amount of Oct4 is required to maintain self-renewal, whereas up or down-regulation of Oct4 leads to directed developmental fates^25^ of ES cells to either PrE or TE, respectively (**Fig. 1a(i)**). On the contrary, Nanog maintains a high expression level in the self-renewing ES cells, but in the other two differentiated states (PrE or TE), its expression level gets down-regulated^26,27^ (**Fig. 1a(ii)**). It seems counterintuitive that in PrE cells, Oct4 expression remains high whereas Nanog maintains a low expression level, as it is well-known experimentally that both Nanog and Oct4 regulate each other’s transcription positively^21,28,29^. Thus, one would expect the Oct4 high level will make the Nanog expression level high as well and vice versa. On the other hand, to maintain the stemness of ES cells, the Nanog expression level must remain at a high state and once differentiated to PrE cells, the Nanog level should go down. This suggests that there are other feedback and feedforward motifs other than mutual positive feedback regulations of Nanog and Oct4 which control these cell fate transitions. However, little is known about the exact dynamical mechanism that finetunes the interactions among the network motifs and orchestrates the cell fate decision-making of ES cells by precisely altering the steady-state expression patterns of Oct4 and Nanog remains elusive.

**Fig. 1:**
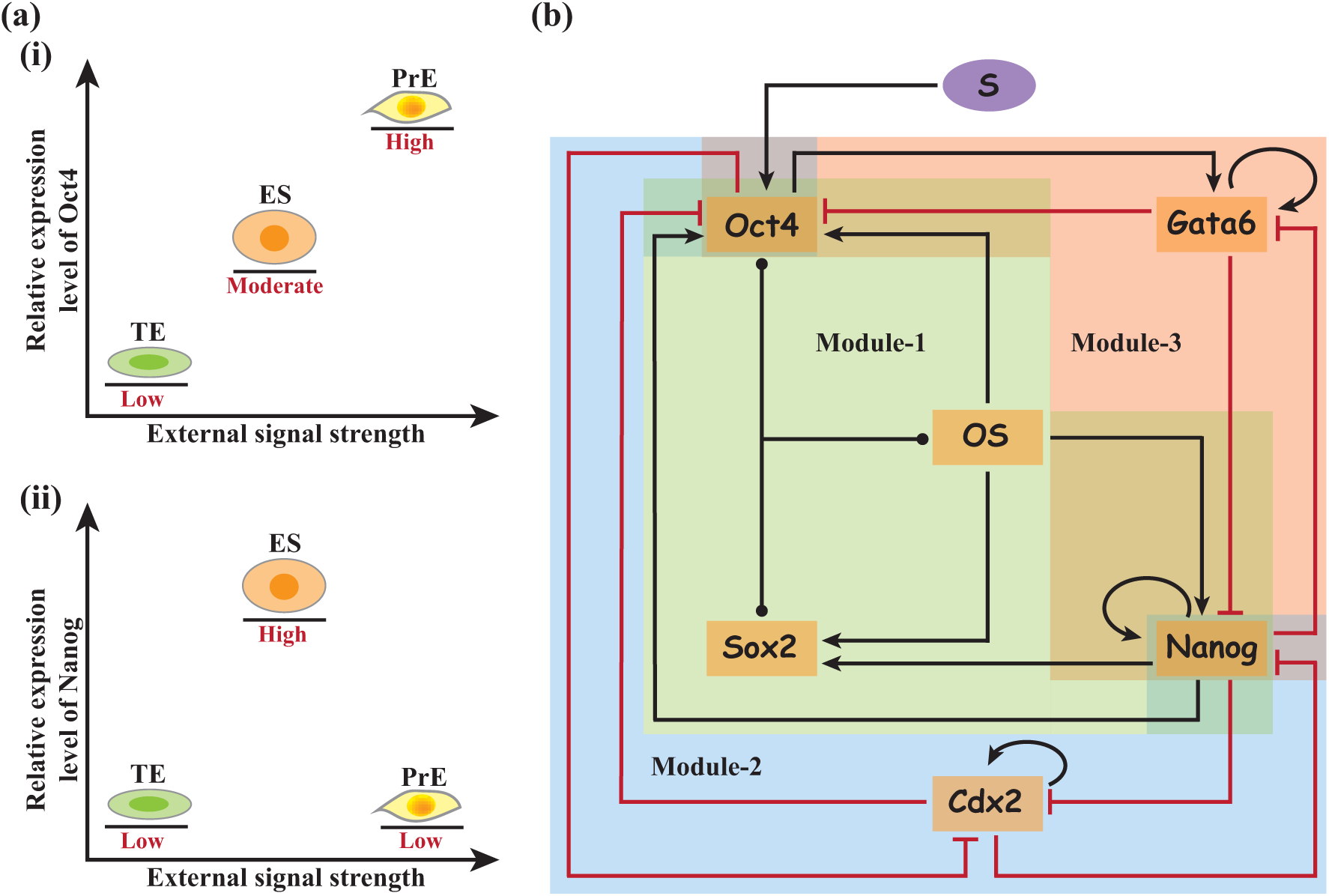
The feedback and feed-forward interaction motifs dictating the cell fate decisions of embryonic stem cells by altering Oct4 and Nanog expression levels. **(a)** A schematic representation of the expression levels of **(i)** Oct4 and **(ii)** Nanog in TE, ES, and PrE cell states. **(b)** Minimalistic molecular interaction network governing the fate commitment events of ES cells. The black solid arrows and red hammer-headed arrows denote biochemical activation and inhibition, respectively.

We must emphasize that in literature, mathematical modelling studies have been employed to understand stem cell differentiation dynamics^21,30–33^. However, most of the models have focused on understanding the pluripotency maintenance of stem cells by considering the core regulatory positive feedback motifs of Oct4-Nanog and Sox2^21,31,34,35^. A few models have looked at the epiblast and PrE specification during embryonic development^7,36^. However, a comprehensive mathematical model that illustrates the possible three different cell fates of ES cells i.e., the stemness maintenance state along with the two differentiated states (TE and PrE) under the influence of varying external signal (S) strength, remains unexplored. Herein, we intend to construct a mathematical model that elucidates the dynamics of stemness maintenance and differentiation towards two extraembryonic lineages of ES cells as per the proposed interaction network shown in **Fig. 1b** where we have considered two key transcription factors Cdx2^37–39^ and Gata6^40,41^ along with the well-established core regulatory network of Oct4-Nanog-Sox2. The deterministic bifurcation analysis of the model suggests that the stemness maintenance of ES cells and its differentiation to either TE or PrE can be explained by two coupled bi-stable switches following stepwise and a mushroom-like steady-state dynamics of Oct4 and Nanog, respectively. The stochastic simulation of the same proposed network (**Fig. 1b**) further predicts the possibility of identifying the existence of the multi-stability of Oct4 as well as Nanog under an appropriate experimental setup. Our mathematical model reconciles the existing experimental observations made in the literature in the context of stemness maintenance of ES cells and differentiation towards the extraembryonic lineages. Importantly, using two-parameter bifurcation analysis, the model identifies the specific role played by various network motifs and qualitatively predicts ways to fine-tune the self-renewal and differentiation dynamics.

## The Model

The developmental dynamics of ES cells are influenced by various regulators and their mutual interactions^4,21^. Among them, Oct4-Sox2 and Nanog form the core regulatory loop in maintaining the stemness of the ES cells, and it is well-established through experiments^17–19^ and computational modeling^21,24,32,34^. Additionally, experimental studies have revealed that transcription factors such as Gata6 and Cdx2 are crucial in determining the differentiation of ES cells towards the extraembryonic lineages^37–41^. The change in the expression pattern of Gata6 in ES cells results in the differentiation of ES cells to PrE cell fate^40,42,43^. A similar alteration in the expression of Cdx2 can cause the differentiation of ES cells to TE cells^27,37^.

Keeping these facts in mind, we have constructed a minimalistic molecular interaction network of these five key transcription factors (**Fig. 1b**) to illustrate the stemness maintenance and differentiation dynamics of ES cells into two extraembryonic lineages under a specific external signal (**S**). This minimalistic interaction network (**Fig. 1b**) can be divided into three modules. Module-1 is comprised of the core regulatory nested positive feedback loop^18,19,44^ among Oct4, Sox2, and Nanog. The transcription factors Oct4 and Sox2 bind to form a heterodimeric complex (OS) that mutually activates all three transcription factors of Module-1. Module-2 represents the TE differentiation. It consists of the mutually antagonistic interaction between Oct4 and Cdx2^37^ and a double negative feedback interaction between Cdx2 and Nanog^45^ along with Cdx2 auto-positive regulation^37,45^. Module-3 is composed of mutual inhibitory interactions between Nanog and Gata6 that govern the PrE fate specification^27,35,46,47^. Further, it is known that Oct4 also activates the transcription of Gata6^41,42,48^. Hence, there exists an incoherent feedforward loop between Oct4, Gata6, and Nanog in Module-3. Additionally, experimental observations suggest the existence of auto-positive feedback interaction present for Gata6^49,50^.

We have translated all the molecular interactions illustrated in **Fig. 1b** into an ordinary differential equation (ODE) based deterministic mathematical model (**Table S1**) that accounts for the evolutionary dynamics of embryonic stem cell development. The related variables are described in **Table S2**. Here, it is important to mention that we have modelled the interactions in a phenomenological manner to explain the stemness maintenance and differentiation dynamics of ES cells by opting for certain kinetic parameters (**Table S3**) such that it allows us to investigate the role of different feedback and feedforward motifs in the context of embryonic stem cell self-renewal and differentiation dynamics.

## Results and Discussions

### Interlinked bi-stable steady-state dynamics of Oct4 and Nanog orchestrate the cell fate decision-making of embryonic stem cells

Niwa et al.^25^ have demonstrated that it is possible to alter the transcription of Oct4 by altering the external signal strength (*S*). Accordingly, we have performed the deterministic bifurcation analysis of our proposed model (**Table S1**) and have found that the steady state level of Oct4 protein undergoes a step-wise bifurcation (**Fig. 2a (i))** as a function of external signal (*S*) due to the existence of two coupled bi-stable switches. On the contrary, Nanog exhibits a distinctive mushroom-shaped steady-state bifurcation (**Fig. 2a (ii))** as a function of *S*. One can interpret such kind of bifurcation pattern of Oct4 and Nanog in the following manner. At lower external signal (*S*) strength, the Oct4 expression level is low and Nanog too maintains a lower expression level. This situation corresponds to the trophectoderm (TE) cellular phenotype. At this range of external signal (*S*) strength, the Cdx2 level is maintained at a high expression level which is a TE marker and the double negative feedback interactions between Oct4-Cdx2 and Nanog-Cdx2 are mainly responsible for generating such a higher expression of Cdx2.

**Fig. 2:**
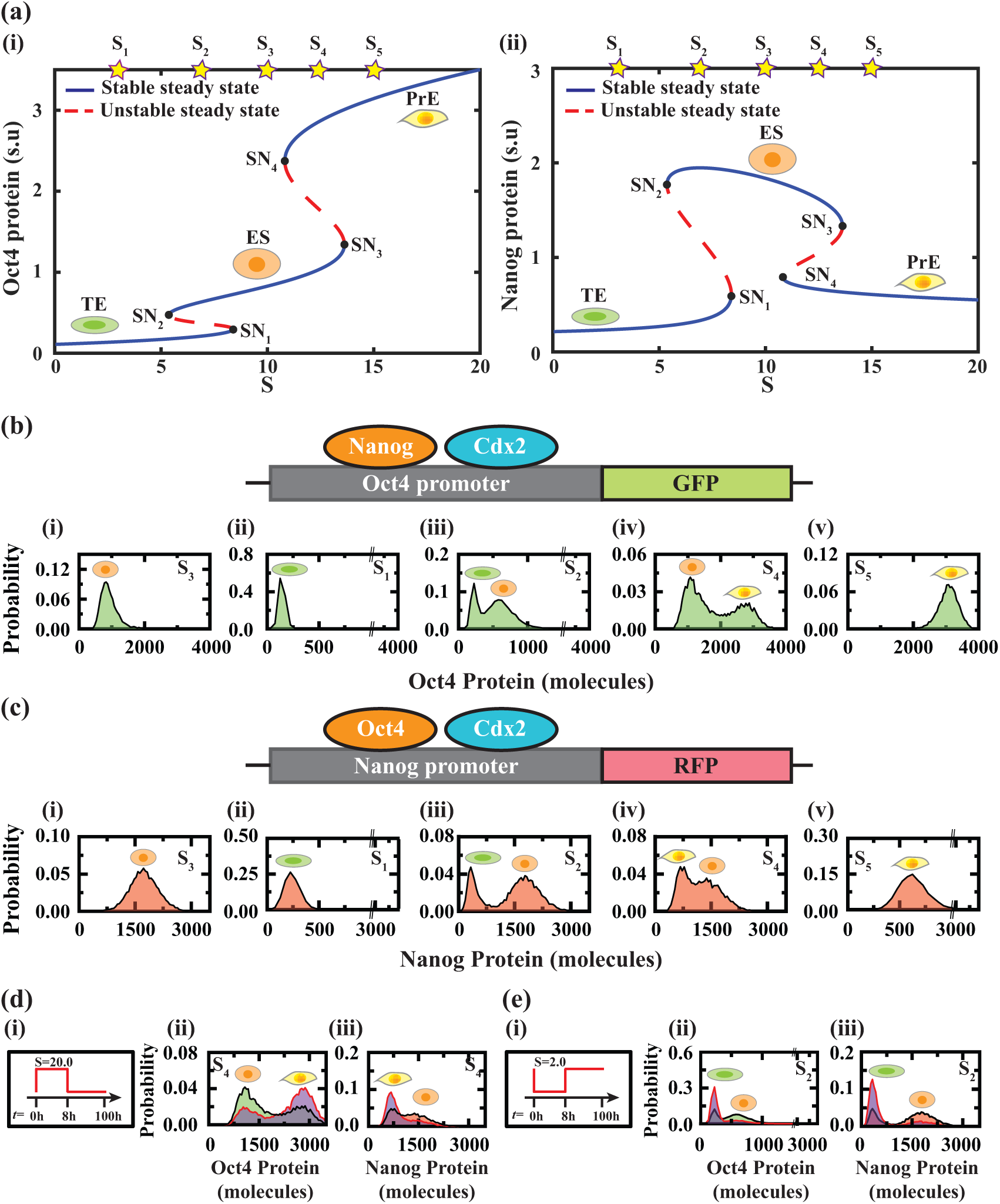
Deterministic and stochastic simulations capture the multi-stable dynamics of Oct4 and Nanog as a function of external signal strength (*S*). **(a)** Deterministic bifurcation diagrams of steady states of (i) Oct4 and (ii) Nanog protein as a function of external signal strength ***S*** (where the saddle nodes appear at ***SN*_1_** = **8.4**, ***SN*_2_** = **5.4**, ***SN*_3_** = **13.6**, and ***SN*_4_** = **10.8**. Distribution of **(b)** Oct4 and **(c)** Nanog proteins at the different external signal strengths, obtained by simulating 10000 stochastic trajectories (or 10^4^ numerical cells) for 100 hours. (***S*_1_** = **3.0**, ***S*_2_** = **7.0**, ***S*_3_** = **10.0**, ***S*_4_** = **12.5**, ***S***_5_ = **15.0**) The distributions of Oct4 protein (at 100 hr) are plotted at 3 different final external signals (***S*_3_** = **10.0**, ***S*_4_** = **12.5**, and ***S*_5_** = **15.0**) after **(d)** a high external signal pulse (***S*** = **20**) is administered numerically (pulse protocol is schematically shown at the top panel) for 8h duration and **(e)** a low external signal pulse (***S*** = **2**) is administered numerically (pulse protocol is schematically shown at the top panel) for 8h duration before simulating at the respective final external signal strengths.

However, as *S* is increased further, the Oct4 level also increases, and the positive feedback interaction between Oct4 and Nanog starts influencing the system. Consequently, the Cdx2 level goes down and beyond a certain threshold *S* level Oct4 attains a moderately higher expression (**Fig. 2a (i))** as Cdx2 switches to a very low level. Nanog steady-state level (**Fig. 2a (ii))** follows the dynamics of the Oct4 steady-state level in this region as Oct4 activates the transcription of Nanog positively. Such a situation where Nanog maintains a high expression level, and Oct4 exhibits a moderate expression level can be mapped with the ES cellular state. Further, an increase in *S* leads to an even higher steady-state level of Oct4 (**Fig. 2a (i))** due to the presence of auto-positive feedback of Oct4 initiated by the Oct4-Sox2 heterodimeric complex and a lower expression of Cdx2 is not able to inhibit Oct4 appreciably. However, in this situation, Nanog reaches a low expression steady state (**Fig. 2a (ii))** by demonstrating a mushroom-like bifurcation. This is caused by the incoherent feed-forward loop involving Oct4-Gata6-Nanog which helps to bring down the Nanog steady state level. Though there is mutual positive feedback between Oct4 and Nanog, the extent of inhibition of Gata6 towards Nanog is overwhelming and it overcomes the Oct4 positive feedback regulation to downregulate the expression level of Nanog. This cellular state where the Oct4 steady state exhibits high expression but Nanog maintains a lower expression can be associated with the PrE fate of ES cells. In this way, our deterministic bifurcation analysis elucidates the differentiation dynamics of ES cells as a function of external signal strength as observed experimentally by Niwa et al.^25^

### Distinct steady-state distributions of Oct4 and Nanog reflect specific cell fate choices

This naturally raises the question, if one follows the Oct4 and Nanog proteins at the single cell by employing suitable single-cell probes, what kind of steady-state distributions of Oct4 and Nanog are expected for the specific cellular fates (i.e., for TE, ES, and PrE states)? Our deterministic model will not be sufficient to answer this important question as it does not account for the fluctuations present in and around the cellular environment. Thus, to get a deeper insight into the developmental dynamics of ES cells, we have used Gillespie’s stochastic simulation algorithm^51^ to account for the inherent stochasticity present in the regulatory network governing the cell fate commitment of ES cells (**Fig. 1b**) under the same parametric conditions (**Table S3**). To verify our hypothesis that two interconnected steady-state bistable switches either in a stepwise (for Oct4) or in a mushroom-like (Nanog) manner as a function of external signal strength controls the differentiation dynamics of ES cells, we have performed stochastic simulations at five different external signal strengths (*S*_1_, *S*_2_, *S*_3_, *S*_4_, and *S*_5_) as depicted in **Fig. 2a-c**). It is important to mention here that in each case, we have started with the initial conditions corresponding to *S*_3_ external signal strength as our bifurcation diagrams suggest that at *S*_3_ signal strength, we will have a pure ES cell population. Furthermore, simulations are performed for 10000 cells at each signal strength, which numerically means simulating 10000 different stochastic trajectories by changing the random number seed during the stochastic simulation.

Indeed, at *S*_3_(S=10) signal strength, our stochastic simulation predicts that the steady-state distributions of Oct4 and Nanog demonstrate uni-modal distributions (**Fig. 2b(i)-c(i)**) with moderate to high and high expression levels of Oct4 and Nanog, respectively. This kind of steady-state distribution can be observed experimentally for such a system by performing flow cytometry experiments which are normally performed to get the expression level of any protein (by tagging the corresponding promoter with a fluorescence protein as shown in **Fig. 2b** and **Fig. 2c** for Oct4 and Nanog, respectively) at the population level under specific signal strengths. The signal strength (*S*) can be systematically varied experimentally in ES cells via the Tc-regulated Oct4 transgene^25^ as demonstrated by Niwa et al.. The steady state distributions for Oct4 and Nanog at *S*_3_ signal strength resembles the ES cellular state. However, once the signal strength is set to a lower value (*S*_1_ = 3), both Oct4 and Nanog show a unimodal distribution with a lower mean expression pattern (**Fig. 2b(ii)-c(ii))**. This happens because, at *S*_1_ signal strength, there exists only one stable steady state at the lower expression level of Oct4 (**Fig. 2a(i)**) and Nanog (**Fig. 2a(ii)**). Thus, our stochastic simulation qualitatively predicts that most of the cells have differentiated into TE cells under this signal strength. Experimental evidence also suggests that TE cells exhibit lower expression for both Oct4 and Nanog^37,45^.

At *S*_2_ = 7.0 signal strength, the system demonstrates bimodal Oct4 and Nanog distributions (**Fig. 2b(iii)-c(iii))** which implies that under this stimulation condition, some of the cells commit differentiation into TE cell fate (low levels of Oct4 and Nanog), and some cells preserve their stemness (moderately high level of Oct4 and high level of Nanog) by maintaining the ES cell state. Interestingly, for higher signal strength (*S*_4_ = 12.0), we again observe bimodal Oct4 and Nanog distributions (**Fig. 2b(iv)-c(iv))**, however, the nature of bimodality differs between Oct4 and Nanog. As we change the signal strength gradually from *S*_3_ to *S*_4_, we observe two different populations of cells, one with a moderately high level of Oct4 and high level of Nanog (ES cells) and the other set of cells demonstrating a high level of Oct4 but a low level of Nanog (PrE cell state). Finally, at a higher value of external signal strength (*S*_5_ = 15.0), most of the cells commit differentiation towards PrE cells exhibiting an unimodal Nanog distribution with a low expression pattern and an unimodal distribution of Oct4 with a higher mean expression level (**Fig. 2b(v)-c(v)**).

Our stochastic simulation results corroborate fairly well with the experimental observations made by Niwa et al.. However, it does not prove our modelling hypothesis that the differentiation dynamics of ES cells are mostly governed by two interconnected bistable steady-state dynamics of Oct4 and Nanog. Thus, we have conceptualized two simulation protocols (**Fig. 2d(i)-e(i)**) to prove our hypothesis which is even possible to validate by performing similar experimental analysis^25^ having suitable fluorescently-tagged single-cell Oct4 and Nanog probes. First, we numerically cultured the cells with high signal strength (*S* = 20) for a short duration (8*h*) (**Fig. 2d(i)**), and for the rest of our simulation time, we simulated the cells with the *S*_4_ signal strength. Our simulation results suggest that under this situation, the probability of expressing high Oct4 (**Fig. 2d(ii)**) and low Nanog (**Fig. 2d(iii)**) levels are indeed becoming more compared to the previous stimulation scenario (**Fig. 2b(iv)-c(iv))**. Thus, under this stimulation protocol, experimentally, we will get a higher extent of PrE cell types. This is because, within the pulse period, the cells that have committed to the stable steady state corresponding to the PrE state, have not been able to return to the steady states corresponding to the ES cell state even when the pulse is taken off. We propose a similar protocol where we start simulating the ES cells with lower signal strength, say at *S* = 2 for a short time duration (8*h*) and then for the rest of the time with the *S*_2_ signal strength, as shown in (**Fig. 2e(i)**). In this scenario, our simulation results predict that we will obtain a higher percentage of TE cells even under *S*_2_ signal strength as demonstrated the respective steady state distributions of Oct4 (**Fig. 2e(ii)**) and Nanog (**Fig. 2e(iii)**). This suggests that the fluctuations present in the system are not enough to propel the cells that are getting stuck at the TE cell state to the ES state. These experiments can establish our hypothesis conclusively.

### Modulating Oct4-Cdx2 double-negative feedback interactions leads to directed differentiation toward TE cell fate

The interaction network (**Fig. 1b**) controlling the differentiation propensities of ES cells consists of several feedback and feedforward network motifs. At this juncture, we have focused on understanding the individual roles played by these motifs in governing the differentiation dynamics of ES cells. First, we have investigated the Oct4-Cdx2 double-negative feedback regulation. Experimentally, one of the routes to affect this feedback motif will be to alter transcription rates of either Oct4 or Cdx2 genes individually. We performed a similar kind of transcription rate variation in our stochastic and deterministic model and followed the steady state distribution and bifurcation pattern of Cdx2 protein (as it is a TE differentiation marker) under different conditions. **Fig. 3a** depicts the distributions of the Cdx2 protein of an ES cell population for the Oct4 downregulation (by reducing the basal transcription rate of Oct4) and Cdx2 overexpression (by increasing the transcription rate of Cdx2) scenarios in comparison to the wild-type situation. **Fig. 3a** foretells that in the wild-type scenario, most of the cells are expressing low levels of Cdx2, while for both the Oct4 downregulation and Cdx2 upregulation situations, more cells are transiting from low level to high levels of Cdx2, which is the signature expression pattern of Cdx2 in TE cells.

**Fig. 3:**
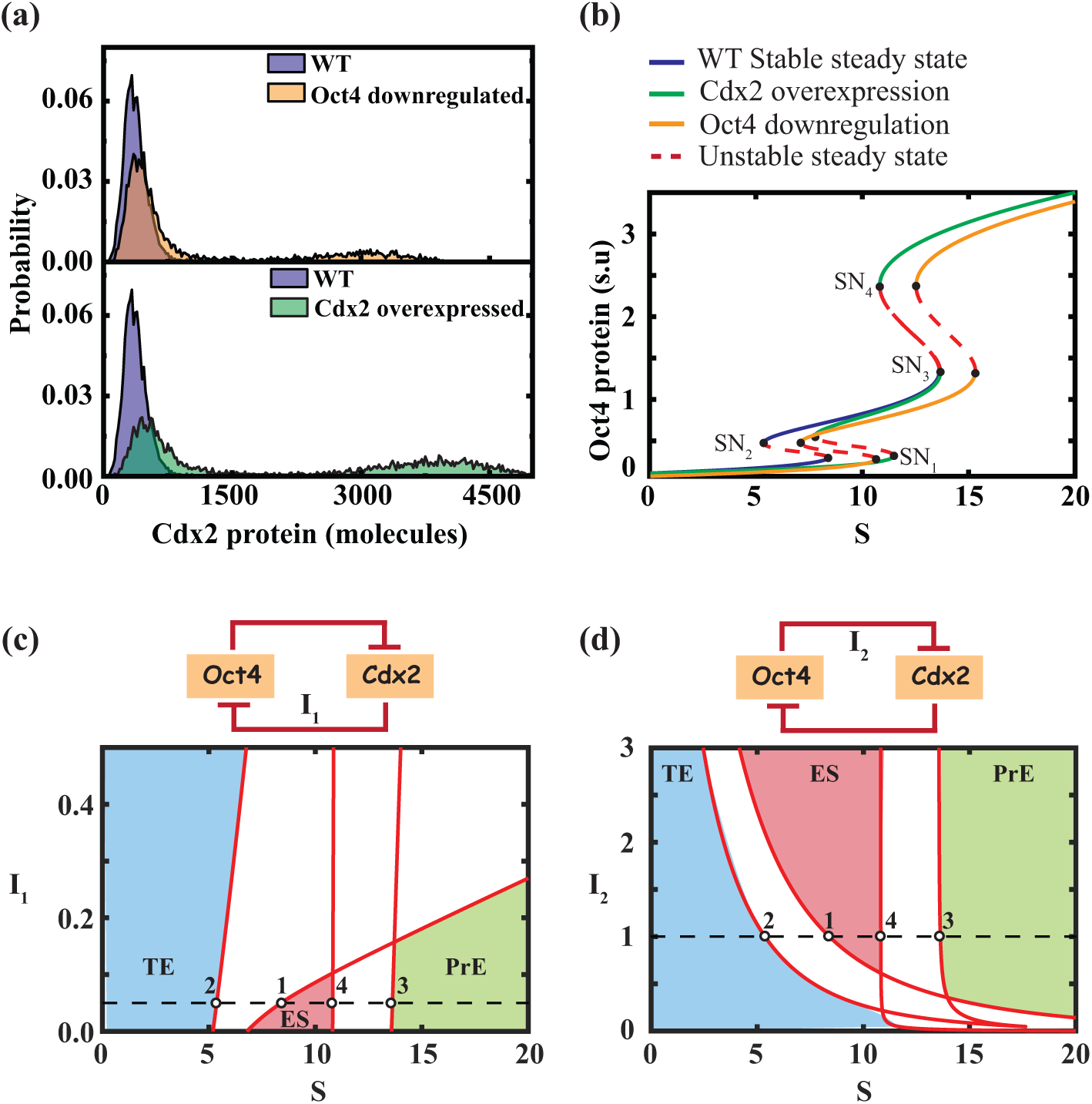
Impact of the double-negative feedback between Oct4 and Cdx2 in regulating the ES cell differentiation propensities. **(a)** Distribution of Cdx2 protein in case of Oct4 downregulation (upper panel) and Cdx2 overexpression (lower panel) in comparison to WT situation. Stochastic simulations are performed for 10000 cells at the external signal strength of ***S*** = **9.0**. **(b)** Bifurcation diagrams of the Oct4 steady-state as a function of ***S*** are plotted for WT, Oct4 downregulation, and Cdx2 overexpression conditions. **(c)** 2p-bifurcation of ***I*_1_** (strength of negative regulation of Cdx2 on Oct4) Vs ***S*** (external signal strength). **(d)** 2p-bifurcation of ***I*_2_** (strength of negative regulation of Oct4 on Cdx2) Vs ***S*** (external signal strength). The dashed line represents the value of the corresponding interaction strength and points 1, 2, 3, and 4 denote the corresponding saddle-node bifurcation points for the WT scenario.

In this regard, it is worth mentioning that Niwa et al. have shown experimentally that either downregulation of Oct4^25^ or overexpression of Cdx2^37^ leads to the differentiation of the ES cells to TE cell types. Our stochastic simulation study performed at the single cell level with Oct4 downregulation and Cdx2 overexpression qualitatively reconciles the experimentally observed population-level behaviour^25,37^ of ES cells. Why more cells are committing differentiation into TE cell fate during Cdx2 overexpression and Oct4 downregulation studies? We get probable reasoning for it by comparing the deterministic bifurcation diagrams (**Fig. 3b**) for the Oct4 as a function of *S* for the wild type, the Oct4-downregulation, and Cdx2-overexpression cases. **Fig. 3b** indicates that for the 1^st^ bi-stable switch, both the saddle nodes (*SN*_1_ and *SN*_2_) shifted towards the right side in the case of the Cdx2 overexpression scenario (**Fig. 3b**) compared to the WT situation (**Fig. 3b**) without altering the saddle-node positions (*SN*_3_ and *SN*_4_) of the 2^nd^ switch. The shift of the *SN*_2_ makes sure that more cells will commit toward TE cells even at the moderate value of the signal strength *S* which for the WT situation only leads to ES cell phenotype. We have a similar observation for the Oct4 downregulation case (stable steady states are denoted by the orange line, **Fig. 3b**) as the 1^st^ switch shifts similarly. However, in this case, the 2^nd^ switch in the Oct4 steady state also shifts towards higher signal strength. This implies that to maintain only the ES-like state or to obtain a pure PrE state, we have to move to relatively higher *S* values, respectively.

To conclusively understand how the double-negative feedback motif between Oct4 and Cdx2 regulates the TE fate specification, we have looked at the two-parameter (2p) bifurcations (**Fig. 3c-d**) by considering the strength of the negative regulations of Cdx2 on Oct4 transcription (*I*_1_) and negative regulation of Oct4 on Cdx2 transcription (*I*_2_), respectively, as the second parameters. The coloured-shaded regions in **Fig. 3c** and **Fig. 3d** represent the different cell fates of ES cells in (*I*_1_, *S*) and (*I*_2_, *S*) parameter spaces, respectively. **Fig. 3c** indicates that if we increase the strength of *I*_1_, the TE cell state will be more preferred in comparison to the ES cell state. Here, with an increase in the negative regulation of Cdx2 (by overexpressing it) on Oct4, the effect of Cdx2 inhibition by Oct4 becomes less. As a result, even at a relatively higher value of an external signal *S*, the ES cells can transit to TE cell fate, which gets reflected in the 1p-bifurcation for the Cdx2 overexpression scenario (**Fig. 3b**). Similarly, one can push the system towards the TE cell fate (**Fig. 3d**) by decreasing the strength of the negative regulation (*I*_2_) of Oct4 (by downregulating it) on Cdx2 transcription. **Fig. 3d** exhibits why with a decrease in *I*_2_, the stemness maintenance region for ES cells decreases, and the propensity to transit at PrE state shifts to higher signal strength.

### Interplay of Cdx2 auto-positive and Cdx2-Nanog double-negative feedbacks govern the ES cells differentiation into TE phenotype

In the proposed differentiation network (**Fig. 1b**), there are two other important network motifs i.e., the Cdx2 auto-positive feedback and double-negative regulation between Nanog-Cdx2, which may play a crucial role in regulating specific features of the ES cell differentiation into TE cell types. To investigate the role of these motifs, First, we look at the time-dependent expression pattern (**Fig. 4a**) of the key transcription factors under the Cdx2 overexpression condition. **Fig. 4a** demonstrates that with time relative mRNA expression of Oct4 and Nanog goes down as Cdx2 negatively regulates both transcription factors. However, the relative changes for Nanog are more compared to that of Oct4. Indeed, Chen et al. have experimentally observed that suppression of Oct4 occurs slower than Nanog^45^. It suggests that Cdx2 regulates both the core regulators Oct4 and Nanog of ES cells in a specific manner. Chen et al. have further observed that the Cdx2 promoter has binding sites for Nanog, Oct4, and Cdx2 itself^45^. To assess the effect of Nanog, Cdx2, and Oct4 on Oct4 transcription, they have co-transfected a Cdx2 reporter plasmid which contains all three binding regions into iRasES cells^45^.

**Fig. 4:**
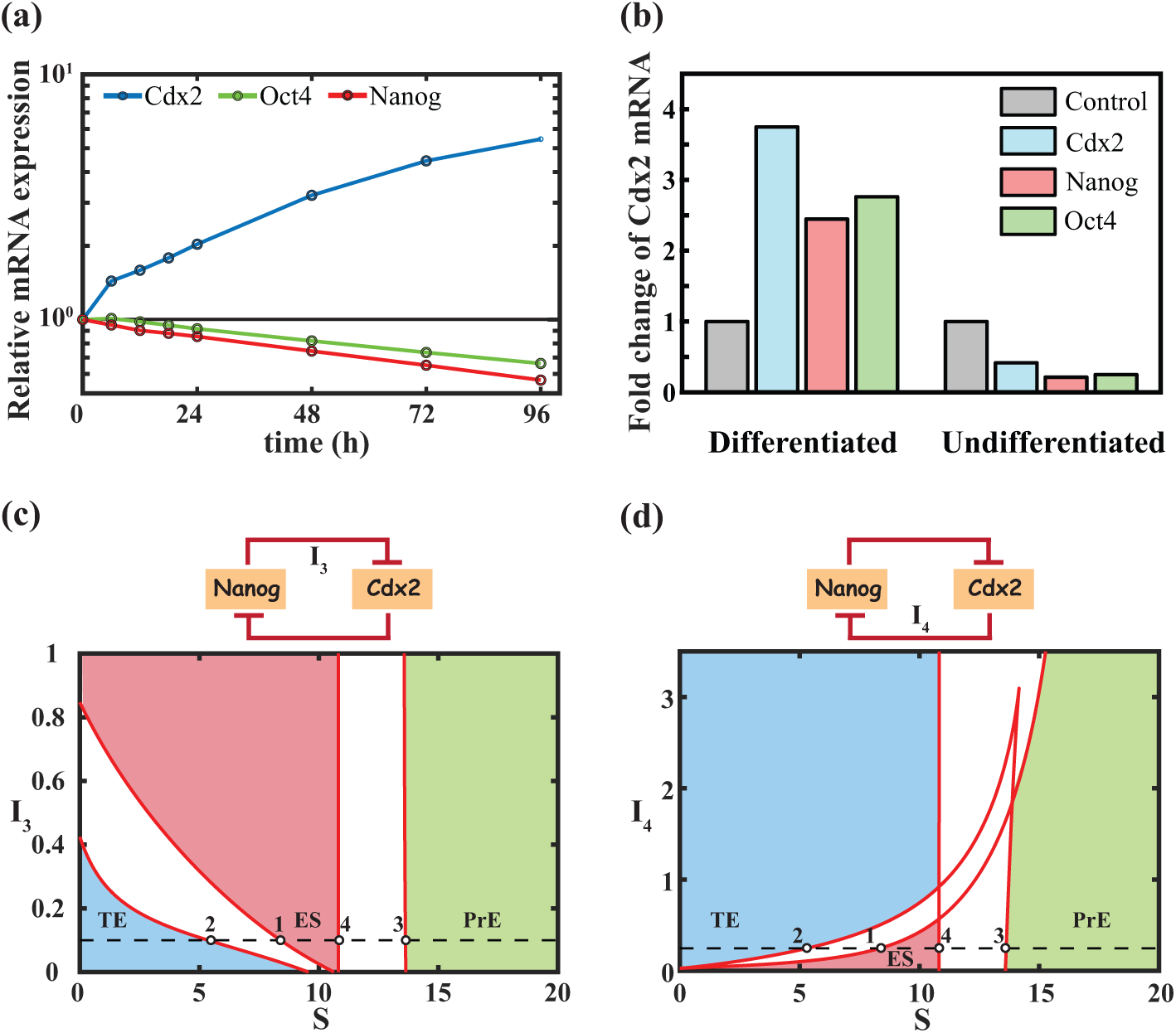
Cross regulation of Nanog-Cdx2 double-negative and Cdx2 auto-positive feedback intricately regulate the TE fate specification of ES cells. **(a)** Temporal dynamics of relative average (10000 numerically simulated cells) mRNA expressions of Oct4, Nanog, and Cdx2 upon Cdx2 overexpression. **(b)** Fold change of Cdx2 mRNA in the case of Cdx2, Oct4, and Nanog overexpression is plotted for differentiated cells (***S*** = **4**. **0**) and undifferentiated cells (***S*** = **10.0**). Here also, the mean value is obtained after performing the simulation for 10000 cells for 96 hours. 2p-bifurcations of **(c)** the strength of negative regulation of Cdx2 on Nanog (***I*_3_**) vs. ***S***, and **(d)** the strength of negative regulation of Nanog on Cdx2 (***I*_4_**) vs. ***S***. The dashed line represents the value of the corresponding interaction strength and points 1, 2, 3, and 4 denote the corresponding saddle-node bifurcation points for the WT scenario (Fig. 2a). The colored shaded regions denote the specified fates of the ES cells while the non-colored regions represent the mixed cellular populations.

In a similar spirit, to study the effect of Nanog and Oct4 in our modelling study, we have numerically overexpressed the Cdx2, Nanog, and Oct4 separately by increasing their transcription rates and measured the fold change of Cdx2 mRNA for two different signal strengths *S* = 4.0 (the differentiated TE cell state region) and *S* = 10.0 (undifferentiated ES cell state region) after 96 hours. **Fig. 4b** shows that in the undifferentiated region, the overexpression of Oct4, Nanog, and Cdx2 suppresses the Cdx2 mRNA expression. However, in the differentiated region, upon overexpression, the Cdx2 mRNA level becomes high, which corroborates with the experimental findings of Chen et al.. This happens because in the undifferentiated region, Cdx2 exhibits lower expression, and the other two pluripotency markers maintain their high levels. As a result, the auto-positive feedback of Cdx2 does not remain fully operational leading to the suppression of Cdx2 mRNA expression. However, in the differentiated region, both Nanog and Oct4 express at lower levels, and the auto-positive feedback of Cdx2 functions effectively. This results in an increase in the Cdx2 level compared to the control one. It is important to mention that in the experiment, the control scenario is represented by the empty vector, which lacks binding sites for Oct4, Nanog, or Cdx2^45^. To replicate a similar control scenario, we simulated the dynamics of Cdx2 mRNA using only the basal synthesis and degradation term, without considering any other regulations (e.g., Oct4 mediated inhibition, Nanog mediated inhibition, and Cdx2 autoregulation). Thus, our modelling studies have found the effectiveness of the feedback motifs in the differentiated and undifferentiated regions of ES cells, which is consistent with the experimental findings^45^.

To further understand the impact of the Cdx2-Nanog double-negative feedback on differentiation dynamics of ES cells, we have performed a 2p-bifurcation analysis concerning the parameters related to the strength of the double-negative feedback interactions (*I*_3_ and *I*_4_). **Fig. 4c** demonstrates that with increasing strength of negative regulation of Nanog on Cdx2 (*I*_3_) at a specific signal strength (*S*), the region of the sole ES cell state increases with a subsequent decrease in the pure TE region which eventually becomes non-existent. Hence, for all the low to moderate signal strengths (between 0-10), with a higher extent of *I*_3_, the sole state is the ES cell state. On the contrary, if we increase the strength of the negative regulation of Cdx2 on Nanog (*I*_4_), the ES cell region becomes shortened, and differentiated TE regions increase (**Fig. 4d**). However, in both scenarios, the PrE region remains unaltered, which suggests that the double-negative feedback regulation of Cdx2 and Nanog does not play any role in the PrE cell fate specification.

### Maintenance of pluripotent state of ES cells by the core regulatory positive feedback motifs formed by Oct4-Sox2 and Nanog

Maintenance of the pluripotent state of the ES cells is extremely important to preserve and use them for therapeutic purposes. Thus, it is imperative to get knowledge about what factors and feedback interactions govern the stemness maintenance of ES cells. Experimental studies indicate that Oct4, Sox2, and Nanog genes play an important role in maintaining the stemness of the ES cells^18,26^ while interacting via interlinked nested positive feedback loops among each other^17,18,44,52^. In this regard, Fong et al. have used target-specific siRNAs to specifically knock down these pluripotency regulators^20^. They have observed that treatment of siRNAs against these transcription factors leads to an increment of TE marker-expressing cells. To mimic such experimental conditions, we have incorporated an additional degradation term in the governing dynamical equation of the three mRNAs (**Table S1**) of Oct4, Sox2, and Nanog. For the wild-type situation, these degradation rates are kept as 0 and are assigned specific values to represent the siRNA knockdown scenarios (**Table S4**).

Numerical simulation concerning the knockdown (KD) studies shows that downregulation of Oct4, Nanog, and Sox2 leads to an increase in the number of cells expressing higher Cdx2 protein (**Fig. 5a**). We have calculated the number of cells expressing higher Cdx2 protein (more than 1500 molecules). The percentage of cells showing higher Cdx2 protein expression (∼43%) almost matches the % of cells showing trophectoderm markers in the experiment performed by Fong et al^20^ (**Fig. 5b**). The reason behind the differentiation of ES cells into TE cells upon knockdown of pluripotency factors lies in the nested positive feedback loops among these three transcription factors. It is well known that Oct4 and Sox2 form a heterodimer complex (OS in our model **Fig. 1b**) that activates all three regulators. Now, knocking down either Oct4 or Sox2 leads to the downregulation of the heterodimeric complex (**Fig. S1**). This downregulation makes the negative regulation of Oct4 on Cdx2 less effective causing more cells expressing higher Cdx2 protein indicative of TE phenotype. The same is true for the Nanog knockdown case, as Nanog positively regulates both Oct4 and Sox2 transcription. Here, we analyze whether it is possible to revive the effect of one pluripotency factor by influencing another pluripotency factor or not. In this regard, Masui et al., have shown that it is possible to rescue Sox2-null mouse ES cells by forced expression of Oct4 through an Oct3/4 transgene^19^.

**Fig. 5:**
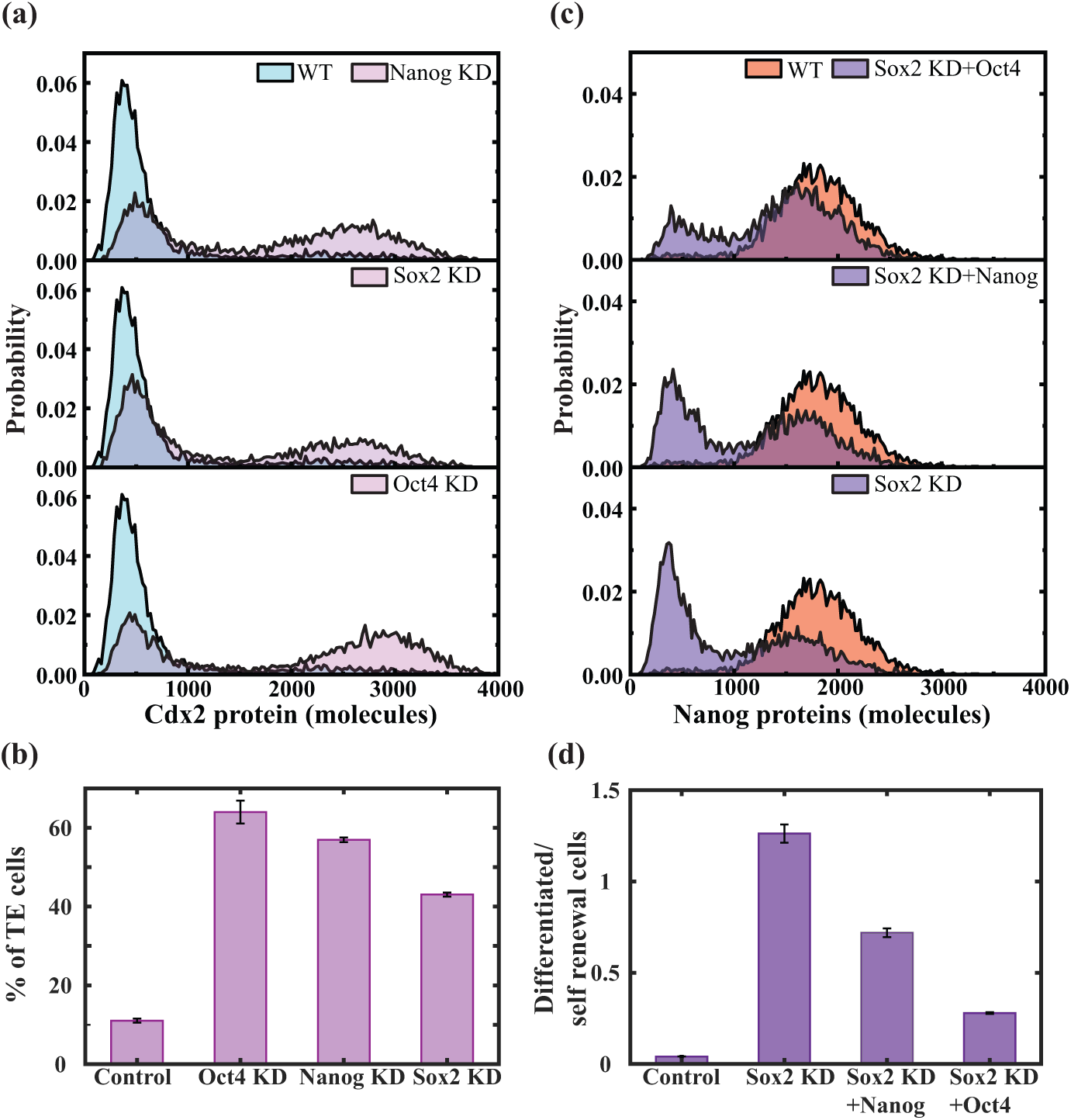
Impact of core pluripotency factors (Oct4-Sox2 and Nanog) in maintaining the stemness of the ES cells. **(a)** Distribution of Cdx2 protein showing knockdown of pluripotency factors leads to TE differentiation. **(b)** Percentage of differentiated TE in the knockdown scenarios. **(c)** Possible to revive pluripotency by inducing Oct4 and Nanog. The distribution of Nanog proteins is plotted for three different scenarios and compared with the wild-type situation. **(d)** The ratio of differentiated TE to self-renewal cells is plotted for the Sox2 KD and the two revived scenarios. The number of TE cells is calculated based on the Cdx2 expression, while the self-renewal cells are calculated based on the expression of Nanog. We have set up a threshold value of Cdx2 protein (1500 molecules), cells expressing Cdx2 protein above that value are considered TE cells. For self-renewing cell counting, the threshold value for Nanog protein is 1000 molecules.

We also try to mimic such culture conditions through our numerical simulations. However, it is not possible in our modelling setup to make a null Sox2 embryo. Instead, we have downregulated the Sox2 expression through our numerical siRNA treatment (**Table S3**) and have investigated whether we can reverse this effect or not. Here, we follow the distribution of Nanog protein to evaluate the self-renewing status of the ES cells. In the wild-type scenario, first, we perform our simulation at S=9.0 signal strength, which is close to the saddle-node point *SN*_1_ and that gives an unimodal distribution for Nanog protein at a higher mean value indicating an ES-like cellular state **Fig. 5c** (WT).

However, in the case of the Sox2 KD scenario, the Nanog distribution turns bimodal, with the appearance of an extra peak in the lower Nanog expressing suggesting the transition of some cells from the ES cell region to the differentiated TE region **Fig. 5c** (Sox2 KD, lower panel). We have simulated 10000 cells to check the revival efficiency to the ES state via over-expressing either Oct4 (**Fig. 5c**, top panel) or Nanog (**Fig. 5c**, middle panel) transcripts under the Sox2 KD situation. **Fig. 5c** predicts that under these overexpressing scenarios, the cells are transiting back to the high Nanog expressing region compared to the Sox2 KD condition. Our calculated ratio of differentiated to self-renewing cells also reflects that it is possible to rescue the self-renewing capacity of ES cells through the forced expression of other pluripotency factors (**Fig. 5d**). This happens because, by overexpressing either Oct4 or Nanog, we can revive the expression of the Oct4-Sox2 enhancer complex, which positively regulates the transcription of all three pluripotency factors. This supports the notion that Oct4-Sox2 and Nanog constitute the core regulatory loop responsible for maintaining the stemness of ES cells. Furthermore, it demonstrates that it is feasible to partially restore the effects produced by any of these core regulators by altering the expression levels of the other core pluripotency factors.

### The incoherent feed-forward loop formed by Oct4-Gata6 and Nanog governs the PrE fate specification

At this point, we further ask how Oct4 controls the development toward the PrE cell specification. In this regard, Frum et al. have shown that Oct4 is required for the development of PrE cells by demonstrating that Oct4-null embryos do not express PrE markers.^42^ We also try to study the role of Oct4 in regulating the PrE specification through our mathematical model. However, in our model, it is difficult to create an Oct4 null situation. Thus, we have studied a similar situation by downregulating the Oct4 and qualitatively compared the results with the experimental findings. We have followed the expression pattern of the Gata6 protein to identify the PrE cell state. **Fig. 6a** (top panel) shows that in the wild-type scenario with *S*_3_ signal strength, most of the cells are maintaining their stemness with a low level of Gata6 expression. However, most of the cells have high Gata6 expression indicating differentiation into PrE cell fate at *S*_5_ signal strength (**Fig. 6a**, bottom panel),. At *S*_4_ signal strength **Fig. 6a** (middle panel), a bistable expression pattern of Gata6 expression portrays that some of the cells have committed differentiation while the rest of the cells maintain their stemness. upon numerical downregulation of Oct4, even at the higher external signal strength (*S*_5_), a bimodal distribution of Gata6 protein (**Fig. 6a**, bottom panel) suggests that some of the cells are now regaining ES cell state and some cells still maintain PrE cell state. However, at *S*_4_, the cells are found to maintain exclusively ES cell state under the Oct4 downregulation condition. This is in stark contrast with the wild-type scenario where a significant proportion of cells have differentiated in the PrE cell state (**Fig. S2**). This shows that Oct4 is highly required for the PrE specification of ES cells, which qualitatively corroborates the experimental findings^42^.

**Fig. 6:**
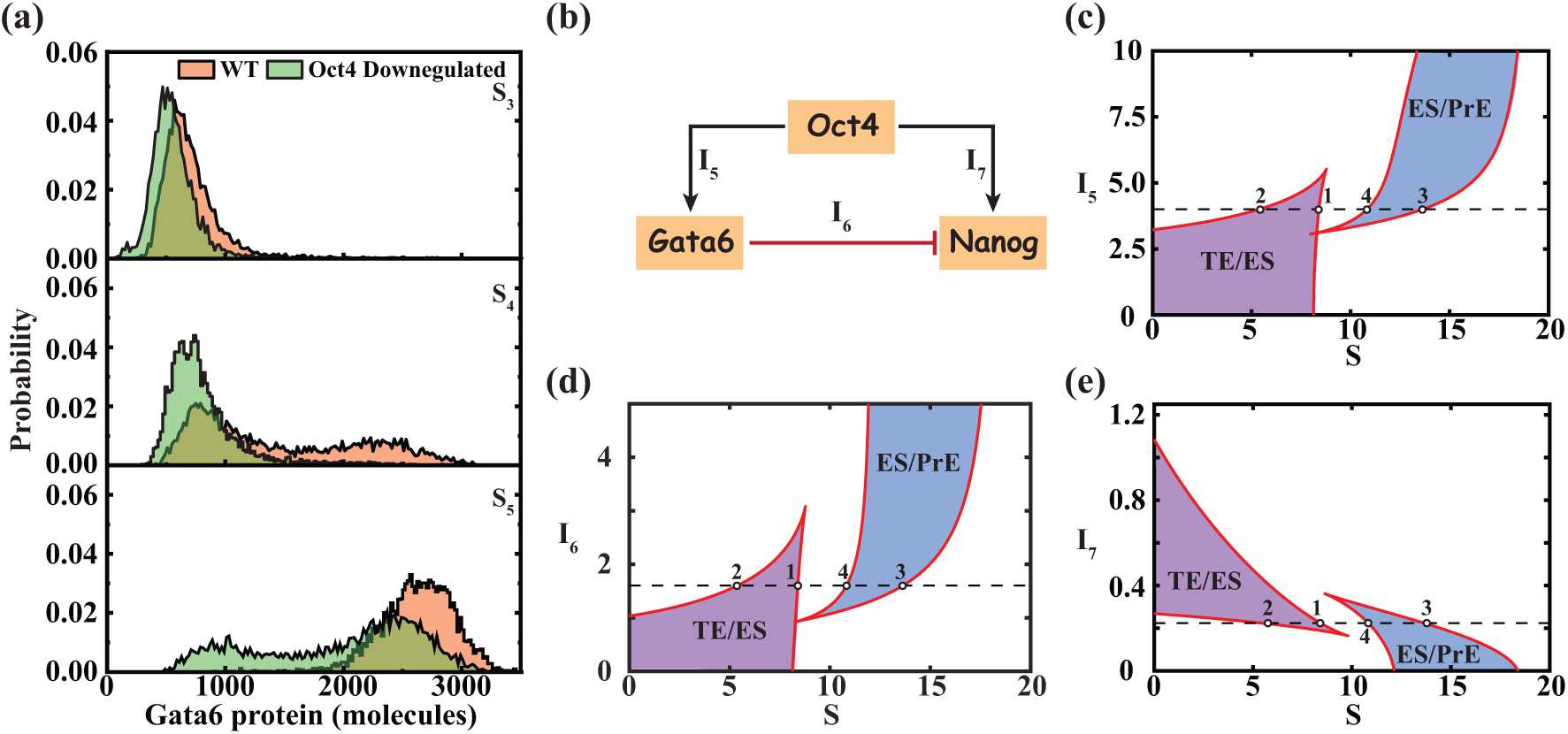
Oct4-mediated incoherence directs PrE cell fate specification of ES cells. **(a)** Distribution of Gata6 protein depicts that downregulation of Oct4 leads to a decrease in the number of cells with high Gata6 expression. Simulations are performed at three different signal strengths (***S*_3_** = **10.0**, ***S*_4_** = **12.5**, ***S*_5_** = **15.0**) for both the WT (***J*_0_** = **0**.**018 *molecules hr*^−1^**) and Oct4 downregulated scenario (***J*_0_** = **0**.**006 *molecules hr*^−1^**). **(b)** Existing incoherent feedforward loop between Oct4-Gata6 and Nanog. ***I*_5_**, ***I*_6_** and ***I*_7_** denote the strength of the corresponding interactions of Oct4-mediated activation of Gata6, Gata6-mediated inhibition of Nanog, and Oct4-mediated activation of Nanog, respectively. 2p-bifurcation analysis between **(c) *I*_5_** Vs ***S***, **(d) *I*_6_** Vs ***S***, and **(e) *I*_7_** Vs ***S*** elucidates the influence of the different interactions on the fate specification of ES cells. (The digits 1, 2, 3, and 4 denote the positions of the saddle nodes (Fig. 2a) under the chosen parametric conditions.)

However, it remains unclear how the downregulation of Oct4 specifically affects the reduction of the PrE cell state population and favors stemness maintenance. This might be because of the existing incoherent feedforward loop between Oct4, Gata6, and Nanog (**Fig. 6b**), where a relative balance between the interaction strengths of Oct4-mediated activation of Gata6 (*I*_5_), Gata6-mediated inhibition of Nanog (*I*_6_) and Oct4-mediated activation of Nanog orchestrate such maintenance of PrE to ES cell state ratio. We have employed a 2p-bifurcation analysis to find how these different interaction strengths regulate the cell fate specifications. **Fig. 6c** demonstrates that with a decrease in *I*_5_(below the dotted line), the *SN*_3_ and *SN*_4_ saddle points will approach each other which makes the bi-stable domain narrower (blue-shaded region). Further decrease in *I*_5_ leads to a cusp point, which means beyond this point, cells will not commit differentiation toward PrE cell fate (**Fig. 6c**). Similar qualitative behavior (**Fig. 6d**) is observed for the Oct4-mediated incoherence of Nanog (*I*_6_).

Interestingly. in one of our previous studies, we showed that by modulating the incoherence in such an incoherent feedforward loop, an Isola kind of bifurcation^53^ can be achieved by altering bi-stable dynamics arising out of a specific set of biological interactions. The molecular interaction network (**Fig. 1b**) also has a similar network architecture, however, **Fig. 6d** demonstrates that it is not possible to get an Isola kind of bifurcation for this differentiation network (**Fig. 1b**). This is because to observe an Isola bifurcation, the *SN*_1_ and *SN*_4_ saddle nodes should annihilate for a specific threshold signal strength (*S*), which is not possible here as the *SN*_3_ and *SN*_4_ points coalesce at the cusp point before the *SN*_4_ saddle-node comes close to *SN*_1_ point (**Fig. 6d**). In hindsight, the appearance of an Isola bifurcation here will be biologically unrealistic, as it will mean that the two different phenotypic cell fates (TE and PrE) are coalescing for specific signal strength. We do not observe an Isola bifurcation here because the two bi-stable switches (producing Mushroom bifurcation of Nanog, **Fig. 1a(ii)**) originate from two different sets of interactions. Here, the 1^st^ bistable switch arises because of the mutual antagonistic nature of Oct4 and Cdx2, and the 2^nd^ bistable switch appears because of the auto-positive feedback interaction of Oct4 through the Oct4-Sox2 heterodimer. However, the reason behind the mushroom-shaped steady-state structure of Nanog is the incoherent feedforward loop (**Fig. 6b** and **Fig. 6c**), which ensures the lower expression of Nanog at higher signal strength even though Oct4 positively regulates Nanog transcription (**Fig. 1a(ii)**). Thus, the incoherent feedforward loop plays an important role in governing the fate specification of PrE cells from the ES cell state (**Fig. 6c**). This is further confirmed when we look at the different steady states in the (*I*_7_, *S*) space (**Fig. 6e**). An increase in *I*_7_ makes the 2^nd^ bistable domain (a blue-shaded region in **Fig. 6e**) narrower, leading to more cells maintaining the self-renewing ES cell state. Thus, our 2p-bifurcation analysis elegantly demonstrates that the incoherent feed-forward loop governs the PrE cell fate specification of ES cells.

### Sensitivity analysis suggests ways to preserve the ES cell state for a wider range of external signal strengths

In the field of stem cell biology, scientists are constantly trying to find ways to either preserve ES cells in a stem cell-like state or figure out strategies to exclusively differentiate ES cells into a specific cell type. Considering these needs, we aim to identify potential ways to alter the dynamics of Oct4 and Nanog to achieve desired cell fate specifications for ES cells. To achieve this, we have conducted a comprehensive sensitivity analysis, considering three specific shaded areas designated by the blue (distance between *SN*_1_ and *SN*_2_ denoted as *D*_1_ = *SN*_1_– *SN*_2_), green (distance between *SN*_3_ and *SN*_4_ denoted as *D*_2_ = *SN*_3_– *SN*_4_), and pink (distance between *SN*_1_ and *SN*_4_ denoted as *D*_14_ = *SN*_1_– *SN*_4_) in **Fig. 7a** as the sensitivity parameters. The *D*_1_ and *D*_2_ represent the bistable regions corresponding to ES-TE cell states and ES-PrE cell states, respectively, while *D*_14_ region denotes the exclusive maintenance of the ES cell state (**Fig 7a**). In our proposed network (**Fig. 1b**), there are six key feedback interactions (**Fig. 7b**) and we have analyzed how these interactions affect *D*_1_, *D*_2_, and *D*_14_ via our sensitivity analysis.

**Fig. 7:**
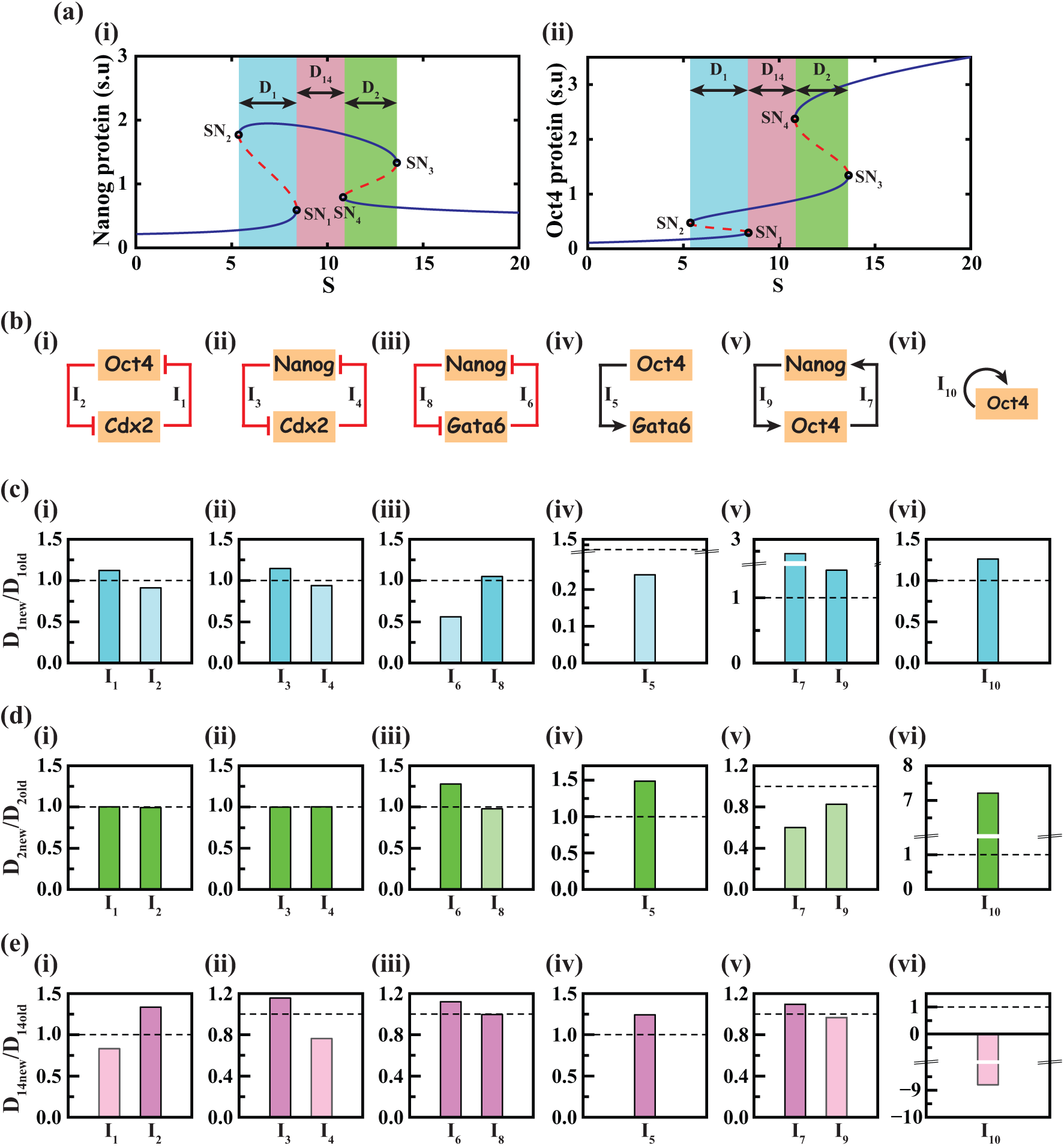
Sensitivity analysis reveals ways to modulate embryonic stem cell developmental dynamics. **(a)** A schematic description of the sensitivity parameters **D_1_** (blue region), **D_2_**(green region) and **D_14_**(pink region), which is defined as *D*_1_ = *SN*_1_ – *SN*_2_, *D*_2_ = *SN*_3_ − *SN*_4_, and *D*_14_ = *SN*_4_ − *SN*_1_ respectively. **(b)** Sensitivity analysis is performed concerning the defined sensitivity parameters for six different feedback loops; (i) Double negative feedback between Oct4 and Cdx2. (ii) Mutual negative feedback between Nanog and Cdx2, (iii) Double negative feedback between Nanog and Gata6, (iv) Oct4-mediated activation of Gata6, (v) Double positive feedback interactions between Oct4 and Nanog, and (vi) auto-positive feedback of Oct4. After performing the sensitivity analysis (increasing each kinetic parameter related to the interaction strength by 20% of the original value (**Table S3**)), we plotted the ratio of **(c)** *D*_1*new*_ to *D*_1*old*_ **(d)** *D*_2*new*_ to *D*_2*old*_ and **(e)** *D*_14*new*_ to *D*_14*old*_. The dark color indicates the value of the ratio is greater than 1 while the lighter color represents the value is less than 1. The negative value of *D*_14*new*_/*D*_14*old*_ indicates that in that scenario, *SN*_4_ comes before the *SN*_1_ point.

**Fig. 7c(i)-(ii)** demonstrates that the first bistable region (*D*_1_) gets marginally modified if any one of the double negative interactions between Oct4-Cdx2 or Nanog-Cdx2 is altered. However, altering the Gata6-mediated inhibition of Nanog (**Fig. 7c(iii)**) and Oct4-induced activation of Gata6 (**Fig. 7c(iv)**) will reduce the bistable *D*_1_ region and favor transition to either ES or TE-like state depending on the signal strength. This observation further confirms that the incoherent feed-forward loop (**Fig. 6b**) plays a pivotal role in either stemness maintenance of ES cells as well as differentiation of ES cells to TE cell type. However, **Fig. 7c(v)** and **Fig. 7c(vi)** depict that by influencing only the mutual activation interactions between Oct4 and Nanog or the auto-activation of Oct4, one can further extend the *D*_1_ bistable region i.e., the region of ES-TE mixed cell state for a wider range of signal strength. This analysis also hints at the sequence of events that underlie the signal strength-mediated differentiation of ES cells. It suggests that diminishing the positive regulations reduces the expression of pluripotency markers (Oct4 and Nanog). This reduction in the expression of Oct4 and Nanog plays a pivotal role in enabling Cdx2 to overcome the inhibitory effects that Oct4 and Nanog typically impose. As a result, the ES cells undergo a transition towards the TE cell state. Hence, sustaining the Nanog-Oct4 positive feedback loop and auto-positive feedback of Oct4 at a moderate level is necessary to uphold the appropriate balance between various phenotypic outcomes.

Sensitivity analysis concerning *D*_2_ region reveals that the Oct4-Cdx2 (**Fig. 7d(i)**) and Nanog-Cdx2 (**Fig. 7d(ii)**) double negative feedback interactions do not affect the ES-PrE coexisting domain. However, sensitivity analysis predicts that the following interactions, namely the Gata6-mediated negative regulation of Nanog (*I*_6_) via Nanog-Gata6 double-negative feedback (**Fig. 7d(iii)**), Oct4-mediated Gata activation (*I*_5_) (**Fig. 7d(iv)**) and Oct4 auto-positive feedback (*I*_10_) regulation (**Fig. 7d(vi)**) will maintain the ES-PrE mixed cellular states for a wider range of signal strength by increasing the 2^nd^ bistable region (*D*_2_). Indeed, enhancing the auto-positive feedback strength of Oct4 (*I*_10_) widens the second bistable switch as the *SN*_4_ point shifts to the left (**Fig. S3(a)**). Interestingly, the fate specification only towards PrE can be primarily governed by increasing the Oct4-Nanog mutual positive feedback regulations (either *I*_7_ or *I*_9)_, (**Fig. 7d(v)**)). Bifurcation analysis confirms that an increase in either of these interaction strengths (*I*_7_ and *I*_9_) results in a narrower *D*_2_ region (**Fig. S3(b)-(c)**). Mechanistically, our sensitivity analysis delineates the dynamical organization of ES to PrE cell state transition in the following manner. Once the Nanog gets over the negative regulation of Gata6 for a threshold level of signal strength, it activates the Oct4 via Nanog-Oct4 positive feedback regulation. A sufficient level of Oct4 activates the Oct4 auto-positive feedback motif to commence the transition to a PrE-like state. That is why the auto-positive feedback of Oct4 influences *D*_2_ region significantly.

Now, we aim to address the question of whether our model can provide insights into achieving an isolated pluripotent ES-like state over a wider range of external signals (*S*). By considering the changes in the sensitivity parameter *D*_14_ (representing the difference between *SN*_4_ and *SN*_1_ saddle points), we have identified a few situations that can lead to greater ES state maintenance (**Fig. 7e**). Specifically our sensitivity analysis predicts that individually increasing the interactions *I*_2_(Oct4-mediated Cdx2 inhibition, **Fig. 7e(i)**), *I*_3_ (Nanog-mediated Cdx2 inhibition, **Fig. 7e(ii)**), *I*_6_(Gata6-mediated inhibition of Nanog (**Fig. 7e(iii)**), *I*_5_ (Oct4-mediated activation of Gata6 (**Fig. 7e(iv)**) and *I*_7_ (Oct4-mediated Nanog activation (**Fig. 7e (v)**) will lead to a greater extent of isolated ES-like cell state. Experimentally, accomplishing any one of these specific modulations will potentially achieve a pluripotent state that remains stable over a wider range of external signals. Thus, our sensitivity analysis provides avenues for controlling and maintaining the desired phenotype of stem cells through targeted adjustments in the feedback interactions highlighted by our model.

### Conclusion

Maintaining the stemness of ES cells and differentiating them precisely to a specific phenotypic cellular state at a desired extent is experimentally quite challenging tasks^11,54^. At the cellular level, these processes are tightly controlled by the varied expression levels of the two key transcription factors, Oct4 and Nanog^18,26^. Often, the phenotypic transitions of the ES cells are shown to be governed by a bistable steady-state expression dynamics of Nanog^30^, where most of the ES cells have a high (ON state) expression of Nanog and a certain fraction of ES cells having a low (OFF state) expression of Nanog will be more prone to differentiate. Intriguingly, the expression level of Oct4 further dictates the cell fate determination of ES cells, by maintaining a low, moderate, and high expression pattern^25^ for TE, ES, and PrE cell types (**Fig. 1a**), respectively. However, it is expected that the Nanog expression level will be low in both the TE and PrE cells but high in the ES cells^26,27^. To explain such kind of complex steady-state dynamics of Nanog and Oct4, by employing mathematical modeling, we hypothesized that the Oct4 and Nanog expression levels in ES cells and the differentiation of ES cells following the molecular interaction network (**Fig. 1b**) are governed by two coupled bi-stable switches arranged in a step-wise (Oct4) and mushroom (Nanog) fashion (**Fig. 2**) as a function of an external signal controlling the transcriptional activation of Oct4.

We have unveiled the complex interplay between two interconnected bistable switches (for both Nanog and Oct4 in a different manner) that may orchestrate the cellular decision-making of ES cells by performing the stochastic analysis of the proposed network (**Fig. 2-6**). We have predicted possible experimental probes that can be engineered to interrogate our hypothesis (**Fig. 2**). Using the steady-state distributions and expression patterns under varied conditions (**Fig. 2-6**), our model we have demonstrated that our simulation results qualitatively corroborate different experimental findings^20,23,25,37,42,45^ on the fate commitments of embryonic stem cells. This exhibits that our initial hypothesis of a stepwise Oct4 and a mushroom-like Nanog steady-state dynamics can indeed control the differentiation dynamics of ES cells as it justifies experimental observations of different kinds. Furthermore, we have unraveled the role of various feedback and feed-forward motifs that are critically controlling the extraembryonic development of ES cells (**Fig. 3-6**). Employing a systematic sensitivity analysis (**Fig. 7**), we have predicted ways to efficiently differentiate the ES cells either to TE or PrE cell types by fine-tuning these feedback interactions. At the same time, it predicts how to maintain the ES cells in a stem-like state for a wider range of signal strength. Importantly, our overall analysis provides a mechanistic interpretation of how the process of differentiation of ES cells to TE and PrE cell types is dynamically organized as a function of signal strength.

In the literature, there are mathematical modeling studies that emphasize the presence of bi-stable switch-like dynamics of Nanog, explaining the occurrence of a bimodal distribution of Nanog^21,24,32,55^. However, to the best of our knowledge, this is one of the first attempts to provide a dynamical interpretation of ES cell development to either TE or PrE cell types based on two interconnected bi-stable switches arranged either in a stepwise (Oct4) or in a mushroom-like (Nanog) steady-state dynamics. Even though right now, it is a hypothesis based on a mathematical and computational study, experimentally, our hypothesis can be verified by performing systematic experimental studies using the reporter constructs (or similar ones) as described in the discussion of **Fig. 2**. The protocol prescribed by us have been followed by experimental groups in the context of cell cycle regulation^56,57^ and epithelial to mesenchymal dynamics^58^. Overall, our study elucidates a highly probable hypothesis that may orchestrate the differentiation dynamics of ES cells to either TE or PrE phenotypes. We have substantiated our hypothesis with bifurcation and stochastic analyses and predicted ways to optimize the self-renewal and differentiation dynamics of embryonic stem cells quantitatively. We hope that this study will generate an interest among experimental biologists to probe the differentiation dynamics of ES cells in an alternative way to generate novel therapeutic strategies.

## Materials and methods

### Deterministic simulations

Our deterministic mathematical model consists of 11 ODEs describing the dynamical evolutions of the mRNAs and proteins regulating the dynamics of extraembryonic development. Deterministic bifurcation analysis of our ODE-based mathematical model (**Table S1**) is performed using XPPAUT (https://sites.pitt.edu/~phase/bard/bardware/xpp/xpp.html). It is worth mentioning that all the mRNAs and proteins are in the scaled unit (s.u) in our deterministic mathematical model. To perform the sensitivity analysis, we have increased the parameter value of the corresponding interactions by 20% one by one and have noted down the positions of the saddle-node points. After that, we performed the analysis on the two bistable regions (TE/ES and ES/PrE) and the sole embryonic stem cell state as mentioned in the main text.

### Stochastic simulations

To incorporate the inherent stochasticity of the cellular environment, we have translated the ODE-based mathematical model into the stochastic model by using chemical master equation formalism. Gillespie’s stochastic simulation algorithm is used to numerically simulate the stochastic mathematical model for incorporating the intrinsic fluctuations of the underlying molecular interactions network (**Fig. 1b**). In our stochastic mathematical model, we have converted the scaled unit of the variables into the number of molecules with the help of a scaling factor (*sf*, mentioned in **Table S3**). The parameter values for the stochastic simulations are given in Table S3, any changes in the parameter values to reproduce the experimental up and downregulations are mentioned in the main text. For the knockdown studies, we have included additional siRNA-mediated degradation terms in the dynamical equations of mRNAs of Oct4, Sox2, and Nanog (**Table S1**). These additional siRNA-mediated degradation terms are kept zero for the wild-type scenarios, and the values of the degradation terms for the knockdown scenarios are mentioned in **Table S4**. It is worth mentioning that we have followed 10000 cells for each numerical simulation, which numerically means picking up different random number seeds for each trajectory.

## Acknowledgments

Thanks are due to UGC for providing the UGC-CSIR-JRF fellowship (Ref. No: 19-06/2016(i)EU-V), IRCC, and the Institute Postdoctoral Fellowship program of IIT Bombay for providing fellowship to AG. This work is supported by the funding agency **SERB, India** (Grant no. **CRG/2019/002640** and Grant no. **MTR/2020/000261**).

